# Neural dynamics of covert habit activation and control in humans

**DOI:** 10.1101/2025.11.12.687990

**Authors:** Eike K. Buabang, Steph Suddell, John P. Grogan, Alice Cox, Nina St. Martin, Kelly R. Donegan, Parnian Rafei, Redmond G. O’Connell, Claire M. Gillan

## Abstract

The neuroscience of habits in humans presumes a competition between two systems: stimulus-response and goal-directed. But lacking strong causal manipulations for inducing habits in humans, researchers have struggled to understand the brain mechanisms supporting the development of stimulus-response associations over time and their interaction with goal-directed processes. To address this, we conducted a preregistered electroencephalography (EEG) study with 14 days of stimulus-response training and used tight temporal controls to isolate the functioning of each system. In line with our preregistered hypothesis, we found that the lateralized readiness potential (LRP) tracked covert activation of habitual responses, detectable on trials where inappropriate habits are overcome and behavior is classed as goal-directed. In exploratory analyses, the strength of this covert habitual activation was associated with the degree of inappropriate habit expression on separate trials where response preparation time was limited. Further, exploratory analyses suggested that frontal midline theta (FMθ) power was selectively elevated when extensively trained habits needed to be overridden, and that this elevation scaled with individual differences in habit expression. These data provide evidence consistent with a dual systems account in which habit expression reflects competition between distinct neural subsystems and offer a new window into the formation and override of habits.

---

Human behavior is thought to reflect a balance between two control systems^1–4^. The goal-directed system selects actions based on expected outcomes and their current value. The stimulus-driven system triggers familiar responses via stimulus–response (S– R) associations, producing habitual behavior that is efficient but inflexible. Habits support the routines of everyday life^5,6^, but some situations demand greater behavioral flexibility and require goal-directed control to override them^7^. Failures to suppress habitual tendencies are a hallmark of compulsivity, including disorders such as obsessive–compulsive disorder (OCD) and disordered eating^8–10^. Mechanistically, inappropriate habit expression is often assumed to occur when the momentary strength of the stimulus–driven process exceeds the engagement of goal-directed control^1^. Yet despite its influence, this account remains quite speculative. Behavioral studies reveal only the outcome of this competition and not the underlying interplay^11^. Most neurophysiological studies have been correlational or limited to causal manipulations on the goal-directed system^12–16^. As a result, there are major gaps in our understanding of how habitual responses are stamped in, activated, expressed, or suppressed.

Repeated training should in theory strengthen habitual tendencies, but demonstrating inappropriate habit expression in humans has been surprisingly challenging, likely because of the overriding influence of goal-directed control^11,17,18^. Recently, however, two paradigms have demonstrated evidence for habits following extensive instrumental training. In the forced-response paradigm, participants learn S–R associations over multiple days and are then tested after a contingency reversal, with response preparation time systematically varied^19^. Extended training leads to more habitual errors when preparation time is limited, consistent with the idea that habitual responses are activated rapidly but are suppressed by goal-directed control when time is sufficient. Using a goal-directed response switching task, it has been shown that overtrained responses produce greater interference, measured as increased response time, when participants must switch to a new goal-directed action following outcome devaluation^20^. Both paradigms converge on the conclusion that extended training strengthens habitual tendencies.

Importantly, however, forced-response measures of habit do not correlate with measures derived from outcome devaluation paradigms^21^, suggesting that these behavioral approaches capture distinct downstream consequences of the balance between stimulus-driven and goal-directed control. Previous EEG studies using outcome devaluation have accordingly identified components related to reward-related stimulus processing and outcome evaluation (P1, P3b, N2, ERN)^22,23^, rather than the motor preparation that is central to stimulus-response accounts of habit. Yet regardless of how habits are assessed behaviorally, the core mechanism posited by dual systems theory is supposed to be the same: a stimulus triggers preparation of a previously trained motor response. A neural measure that directly indexes this motor preparation should therefore capture the strength of the habitual association independently of the specific behavioral paradigm used to probe it.

The lateralized readiness potential (LRP) offers precisely this as it reflects effector-specific motor preparation over the contralateral motor cortex, providing a continuous readout of which response is being prepared even when that response is not ultimately executed^24^. Here, we leveraged this to address the gap between behavioral and neural accounts of habit. Participants completed a within-person study comprising 1-day vs 14-day training of four S–R associations. After each training condition, there was a contingency reversal in which the responses for two stimuli were remapped across hands, resulting in the correct response being mapped to one hand and the habitual, originally learned response to the opposite hand. This allowed the LRP to track the covert preparation of the habitual response even on trials where it was successfully overridden. We indexed goal-directed control with frontal midline theta (FMθ) power, a marker that increases with demands on cognitive regulation and control^25–28^. Participants were tested with either short or long response preparation times, where the former was designed to reveal habits behaviorally, and the latter designed to measure EEG signatures unconfounded by behavioral differences.

In line with our pre-registration, we found that the LRP indexed covert activation of the habitual response when participants had been overtrained even on trials where participants had ample time to prepare their response and selected the correct action. Exploratory analyses revealed that the strength of this habitual EEG signature was associated with behavior (i.e., habit errors) on a separate set of short response preparation trials. Exploratory analyses of FMθ suggested that reactive goal-directed control was selectively elevated when extensively trained habits needed to be overridden. This causal result was corroborated by correlational data showing that this elevation scaled with individual differences in habit expression. Together, these findings provide new evidence consistent with the view that the expression of habitual behavior arises from competition between distinct neural subsystems. In doing so, our study offers a novel platform for observing the interplay between stimulus-driven and goal-directed control systems, which can be applied to better understanding the mechanisms leading to compulsive psychopathology and other maladaptive behaviors^28^.

## Results

### Behavioral evidence for stimulus-driven and goal-directed control

In a preregistered within-person crossover design, participants completed minimal (1-day) and extensive (14-day) training on a habit task^19^, with the order of training conditions counterbalanced across participants (Fig. 1a). During training, participants first learned an original mapping consisting of four arbitrary symbol-keypress associations via trial-and-error (Fig. 1b,c). In both training conditions, initial acquisition required participants to reach a learning criterion (i.e., five consecutive correct responses per stimulus). This criterion training was the only training provided in the minimal condition and the starting point for the extensive condition, where participants then continued training across 12 remote sessions, with in-lab training sessions on both Day 1 and Day 14. Following the minimal and extensive training period, participants learned a revised mapping to criterion, where two S-R associations were remapped and two remained consistent (Fig. 1d). This remapping enabled the measurement of habit errors, defined as responses in line with the original but now-incorrect mapping. In the subsequent forced-response test (Fig. 1e), participants were required to synchronize their response with a sequence of three tones. Stimuli appeared either 400 ms (short preparation) or 1100 ms (long preparation) before the final tone that served as response cue, thereby manipulating the time allowed for action selection. Prior research has shown that habits are more evident on trials with short preparation, where participants theoretically do not have sufficient time to engage goal-directed resources^19^. The final sample consisted of 30 participants (*M*_age_ = 21.3 years, *SD* = 3.1; 19 female, 9 male, 2 non-binary; see Methods for exclusions). For EEG analyses, additional exclusions were applied during preprocessing, resulting in final sample sizes of *n* = 23 for the LRP analyses of the forced-response test, *n* = 28 for the FMθ power analyses of the forced-response test, and *n* = 29 for the training analyses. Preregistration, data, analysis code, and supplemental materials are available on the [Open Science Framework](https://osf.io/fm8q7/overview?view_only=5a02494967ad451eb6162dd566dcc8df).

**Fig. 1:**
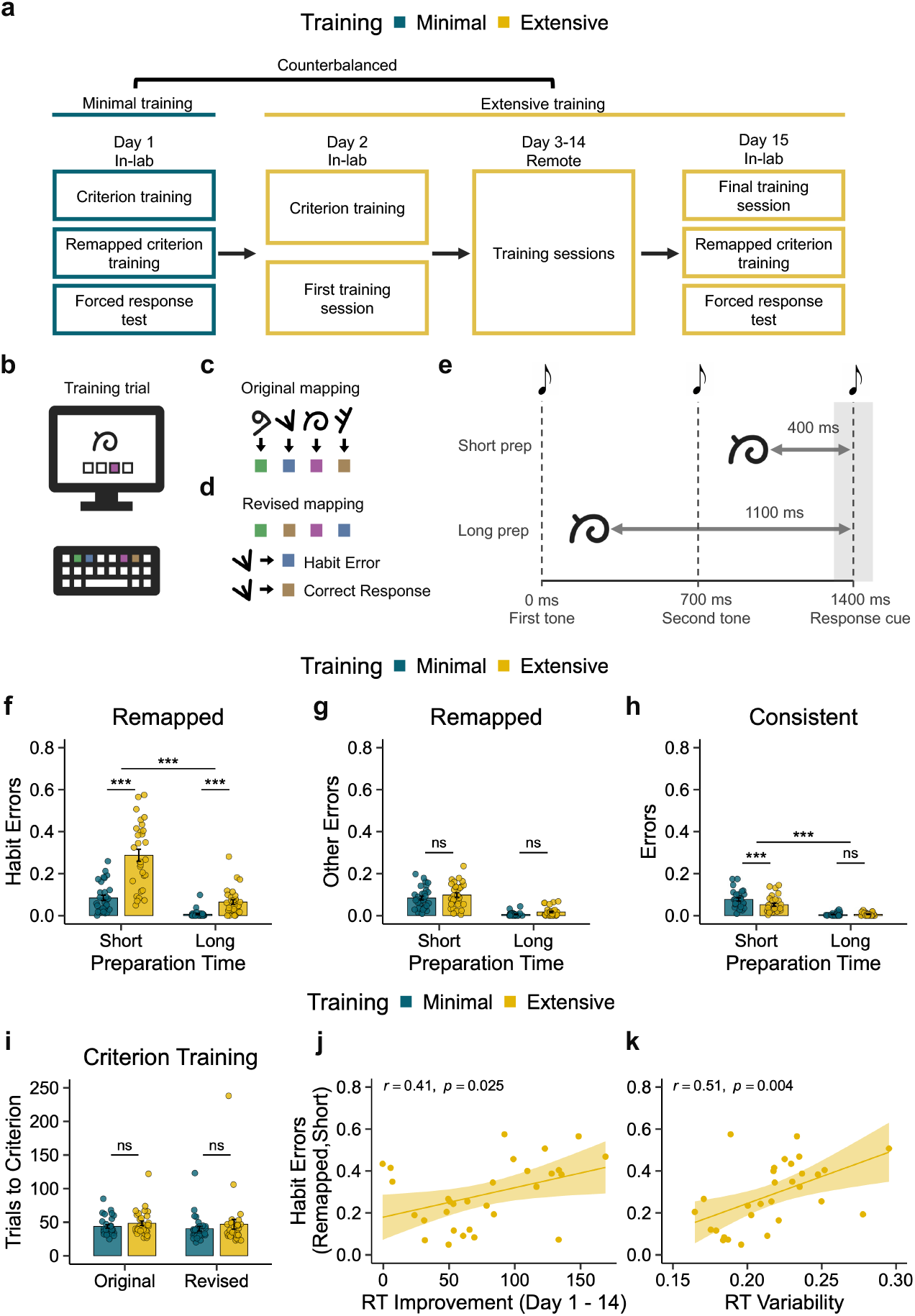
Task design, training effects, and individual differences in habit expression. **a**, Overview of the within-participant crossover design. Participants completed both minimal (teal) and extensive (yellow) training conditions in counterbalanced order. The minimal condition involved a single in-lab session; the extensive condition included ∼9,000 trials over 14 days. **b**, Example training trial. Participants learned to select the correct keypress in response to arbitrary symbols. **c**, Participants first learned an original mapping consisting of 4 symbol–keypress associations. **d**, A revised mapping was then introduced, allowing detection of habit errors, defined as responses consistent with the original but now-incorrect association. **e**, During the forced-response test stimuli were presented either 400 ms (short preparation) or 1100 ms (long preparation) before the auditory response cue, limiting or allowing time for goal-directed selection. **f**, The proportion of habit errors increased significantly after extensive training, particularly under short preparation (Training × Preparation Time: *F*_1,29_ = 29.32, *p* < 0.001; short: *t*_29_ = 7.49, *p* < 0.001; long: *t*_29_ = 5.20, *p* < 0.001). **g**, Other errors increased under short preparation time (*F*_1,29_ = 113.65, *p* < 0.001) but did not vary with training duration (all *F*s < 3.74). **h**, The proportion of errors on consistent trials was lower after extensive than minimal training under short preparation (*F*_1,29_ = 18.89, *p* < 0.001; short: *t*_29_ = 3.91, *p* < 0.001). **i**, The number of trials to reach criterion did not differ between training conditions or mapping (all *F*s < 1.81). **j**, Greater RT improvement over the 14-day training period was associated with more habit errors at test (*r* = 0.41, *p* = 0.025). **k**, RT variability across training, quantified as the coefficient of variation (standard deviation / mean RT), was also positively correlated with habit errors (*r* = 0.51, *p* = 0.004). Dots reflect individual participants. Bars show mean ±1 SEM; shaded regions represent 95% CIs. \*\*\**p* < 0.001; ns = not significant.

As predicted (H_1_), extensive training increased the likelihood of inappropriate habit expression during the forced-response test, particularly when response preparation time was limited. For remapped trials, an ANOVA on the proportion of habit errors revealed a significant interaction between Training and Preparation Time (*F*_1,29_ = 29.32, *p* < 0.001; Fig. 1f). Habit errors were more frequent after extensive (0.29 ± 0.16) compared to minimal training (0.08 ± 0.07) on short preparation trials (*t*_29_ = 7.49, *p* < 0.001). The difference between extensive and minimal training was smaller but remained significant on long preparation trials (0.06 ± 0.06 vs. 0.01 ± 0.02; *t*_29_ = 5.20, *p* < 0.001). Main effects were also observed for both Training (*F*_1,29_ = 68.31, *p* < 0.001) and Preparation Time (*F*_1,29_ = 98.94, *p* < 0.001), reflecting overall increases in habit errors following extensive vs. minimal training and on short vs. long preparation trials, respectively. These results replicate prior findings^19,28–31^and support the conclusion that extensive training strengthens habitual response tendencies, which exert strong influence when time constraints limit goal-directed control.

To confirm that our findings specifically relate to habit errors, we also analyzed other errors on remapped trials, defined as responses that matched neither the revised nor the original mappings. Other errors increased under time pressure (*F*_1,29_ = 113.65, *p* < 0.001; Fig. 1g), suggesting a general speed-accuracy trade-off, but importantly there was no significant effect involving training duration (*F* < 3.74).

We next explored performance on consistent trials, where mappings remained unchanged. An ANOVA revealed a significant interaction between Training and Preparation Time (*F*_1,29_ = 18.89, *p* < 0.001; Fig. 1h). The proportion of errors was significantly lower after extensive (0.05 ± 0.04) compared to minimal training (0.08 ± 0.04) on short preparation trials (*t*_29_ = 3.91, *p* < 0.001). Thus, the pattern was the opposite of that for remapped trials, indicating that extensive training promotes rapid correct responding for stable S-R mappings. There was no significant difference on long preparation trials (*t*_29_ = -1.23, *p* = 0.227). Together, the results from remapped and consistent trials indicate that repetition strengthens habitual tendencies and their expression under time pressure, but whether they are appropriate or inappropriate depends on the context.

We also confirmed that the results during the forced-response test were not due to differences in learning. There was no significant difference in the number of trials to reach the criterion and learn the original and revised mappings for both training conditions (all *F*s < 1.81; Fig. 1i), indicating that the acquisition of the S-R associations was similar for the two training conditions.

Next, we explored how habit errors related to performance over the 14-day training period. Although reaction times (RTs) reduced overall across days, signs of fatigue emerged within sessions, as reflected in increasing RT variability and error rates (see Extended data). Exploratory analyses further revealed that participants who showed greater improvements in RTs from Day 1 to Day 14 made more habit errors at test (i.e., on remapped trials with short preparation time after extensive training: *r* = 0.41, *p* = 0.025; Fig. 1j), suggesting greater automatization of responses. Additionally, these habit errors were positively correlated with RT variability across training (*r* = 0.51, *p* = 0.004; Fig. 1k), quantified as the coefficient of variation (standard deviation divided by mean RT).

### Covert motor preparation reveals activation of habitual responses

To assess the neural signature of the stimulus-driven system, we analyzed the LRP, a well-established index of effector-specific motor preparation, derived from EEG activity over motor cortex sites contralateral versus ipsilateral to the responding hand^24^. In our study, the LRP reflects the difference in motor preparation between the correct and incorrect hands. Because remapping required switching two S–R associations across hands, the design enabled us to directly assess motor preparation for the correct response hand relative to the originally trained but now incorrect response. We predicted that the LRP would reveal preparation of the habitual response even on trials where participants ultimately selected the correct action. To this end, we focused on long-preparation trials of the forced-response test, where participants had ample time to select the appropriate response, and restricted analyses to correct trials to isolate covert activation unconfounded by overt habit errors.

We first conducted the preregistered 2×2 repeated measures ANOVA on mean LRP amplitude across the 150–800 ms window, testing the interaction between Training and Mapping. As predicted (H_2_), this revealed a significant Training × Mapping interaction (*F*(1,22) = 4.42, *p* = .047; Fig. 2a), with no significant main effects (Mapping: *F*(1,22) = 0.06, *p* = .807; Training: *F*(1,22) = 0.04, *p* = .842). The interaction reflected a crossover pattern.

**Fig. 2:**
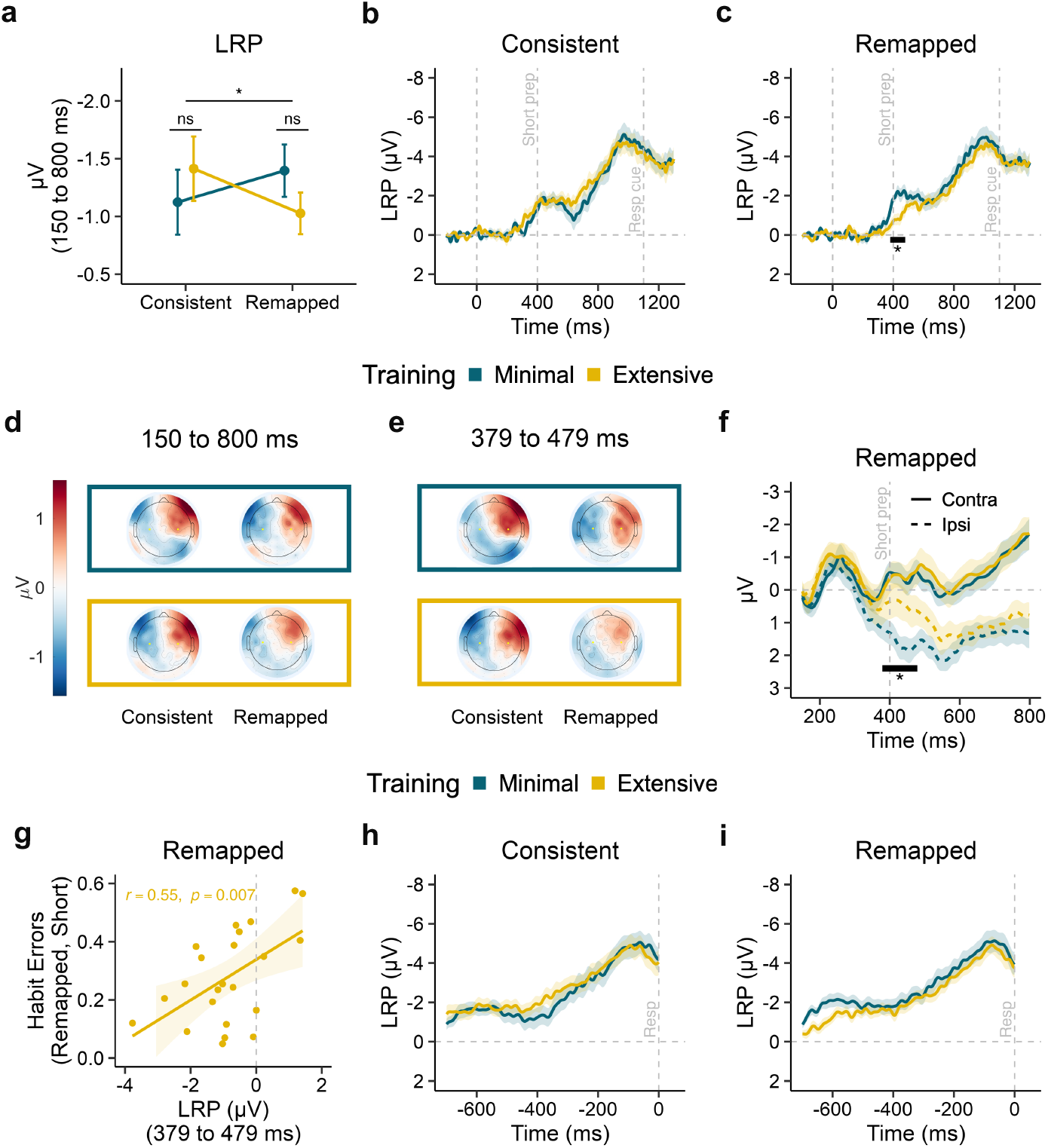
LRP indexes covert activation of habitual responses and scales with individual habit strength. **a**, Mean LRP amplitude (150–800 ms) showed a significant Training × Mapping crossover interaction (*F*(1,22) = 4.42, *p* = .047). **b**, Grand-average stimulus-locked LRPs on consistent trials following minimal and extensive training. No significant clusters were observed. **c**, Grand-average stimulus-locked LRPs on remapped trials. A significant training effect emerged between 379 and 479 ms (black bar; cluster-based *p* = .038), with reduced LRP amplitudes after extensive training, consistent with covert activation of the habitual response. **d**, Scalp topographies of the LRP during the 150–800 ms window for each condition. Yellow dots indicate LRP electrode sites (C3, C4). **e**, Contralateral and ipsilateral activity on remapped trials. A significant Training × Hemisphere interaction emerged between 379 and 479 ms (cluster-based *p* = .037), driven by increased ipsilateral (habitual response hand) activation after extensive training (*t*(22) = 2.25, *p* = .035), with no change in contralateral activation (*t*(22) = −0.04, *p* = .965). **f**, Scalp topographies during the 379–479 ms cluster window. **g**, LRP amplitude on remapped trials after extensive training (379– 479 ms) significantly predicted habit errors (*β* = 0.070, *p* = .007). **h**, Grand-average response-locked LRPs (−700 to 0 ms) for remapped trials. **i**, Response-locked LRPs for consistent trials. No significant effects were observed in the response-locked analysis. Shaded regions represent ± 1 SEM. \**p* < .05; \*\**p* < .01; ns = not significant.

After extensive training, LRP amplitudes were numerically larger on consistent trials (*M* = −1.41 µV, *SD* = 1.34) than after minimal training (*M* = −1.12 µV, *SD* = 1.35), whereas the pattern reversed on remapped trials (extensive: *M* = −1.03 µV, *SD* = 0.87; minimal: *M* = −1.40 µV, *SD* = 1.08). Follow-up paired t-tests within each mapping did not reach significance individually (consistent: *t*(22) = −1.26, *p* = .219; remapped: *t*(22) = 1.37, *p* = .185), indicating that the interaction was driven by the opposing pattern across both trial types.

Within the full 150–800 ms window, we hypothesized that habitual activation would specifically manifest as an early Gratton dip, a brief polarity reversal, from 150–300 ms, measured as the positive area under the LRP curve. However, this was not supported (interaction: *F*(1,22) = 0.04, *p* = .842; all *F*s < 0.17, all *p*s > .68). To understand what time-window was driving the significant interaction, we performed an exploratory cluster-based permutation analysis on the stimulus-locked LRP time series (150 to 800 ms) for each mapping separately. No significant clusters were observed for consistent trials (Fig. 2b). But on remapped trials, LRP amplitudes were significantly reduced following extensive compared to minimal training between 379 and 479 ms (cluster-based *p* = .038; Fig. 2c). This reduction in LRP amplitude reflects a relative shift in motor preparation toward the habitual response hand. Later in the time series than we anticipated, the timing of this effect coincides with the onset of the short-preparation response cue at 400 ms. This appears to suggest that habitual response activation on long preparation trials occurs at the same time as participants typically exhibit habits on short preparation trials. Scalp topographies showed the expected lateralization pattern (Fig. 2d,e).

Because the LRP is a relative measure, reduced amplitudes on remapped trials following extensive training could reflect less activation for the correct response (contralateral activity), increased activation for the habitual response (ipsilateral activity), or both. To disentangle these contributions, we repeated the cluster-based permutation analysis on the stimulus-locked LRP time series (150 to 800 ms), this time evaluating the interaction between Training and Hemisphere (contralateral vs. ipsilateral). This revealed a significant interaction between 379 and 479 ms (cluster-based *p* = .037; Fig. 2f), coinciding with the time window identified in the main LRP analysis. Follow-up paired t-tests within this window showed that ipsilateral activation was significantly greater after extensive compared to minimal training (*t*(22) = 2.25, *p* = .035), whereas contralateral activation did not differ (*t*(22) = −0.04, *p* = .965). This indicates that the LRP reduction on remapped trials was driven by increased motor preparation for the habitual response hand, rather than decreased preparation for the correct response.

We next tested whether covert habit activation related to behavioral habit expression. We regressed the proportion of habit errors after extensive training (remapped, short) on LRP amplitude from remapped trials after extensive training. Following our pre-registration (H_3_), we tested this on the entire 150–800 ms window, but this was not significant (*β* = 0.031, *SE* = 0.040, *t*(21) = 0.78, *p* = .446). However, this is perhaps unsurprising, as our previous analysis showed that covert habit activity was evident in a much tighter window of 379–479 ms. We thus performed an exploratory analysis using the crucial habit window and found that LRP amplitude significantly predicted habit errors (*β* = 0.070, *SE* = 0.023, *t*(21) = 3.00, *p* = .007; Fig. 2g). Specifically, individuals with reduced LRP amplitudes, reflecting stronger covert habitual activation, made more habit errors on our crucial test trials (i.e. when they were under time pressure). On consistent trials, the pattern reversed. Larger LRP amplitudes, reflecting stronger correct-response preparation, were associated with more habit errors (*β* = −0.040, *SE* = 0.019, *t*(21) = −2.14, *p* = .044). Together, these results suggest that a common habit mechanism may drive both enhanced correct-response preparation on consistent trials and covert habitual activation on remapped trials, with both scaling with individual differences in behavioral habit expression.

Finally, we complemented the stimulus-locked analysis with a preregistered 2×2 repeated measures ANOVA on mean response-locked LRP amplitude (−700 to 0 ms), testing the interaction between Training and Mapping. No significant effects were observed (interaction: *F*(1,22) = 2.07, *p* = .165; all *F*s < 2.07, all *p*s > .165; Fig. 2h,i). This is consistent with participants having successfully overridden habitual motor preparation by the time of response execution.

### Frontal midline theta reflects goal-directed control demands

To assess goal-directed control, we analyzed FMθ power, a well-established index of cognitive control, implicated in conflict monitoring and the resolution of competing response tendencies^25–27^. We predicted that FMθ power would be selectively elevated on remapped trials after extensive training, reflecting increased recruitment of goal-directed control to override habitual response tendencies. Our preregistration focused on the 0 to 500 ms post-stimulus window, which includes the emergence of the covert habit activation. As with the LRP analyses, we focused on long-preparation, correct trials to isolate control engagement unconfounded by overt habit errors.

Contrary to our prediction (H_4_), the 2×2 repeated measures ANOVA on mean FMθ power during the 0–500 ms window revealed no significant Training × Mapping interaction (*F*(1,27) = 0.00, *p* = .983; Fig. 3a,b), indicating no selective increase in FMθ on remapped trials after extensive training. However, significant main effects were observed. FMθ power was higher on remapped than consistent trials (*F*(1,27) = 4.73, *p* = .039), and higher after extensive than minimal training (*F*(1,27) = 6.64, *p* = .016). These main effects suggest that FMθ increases with both mapping conflict and training history, though not in the condition-specific pattern we predicted.

**Fig. 3:**
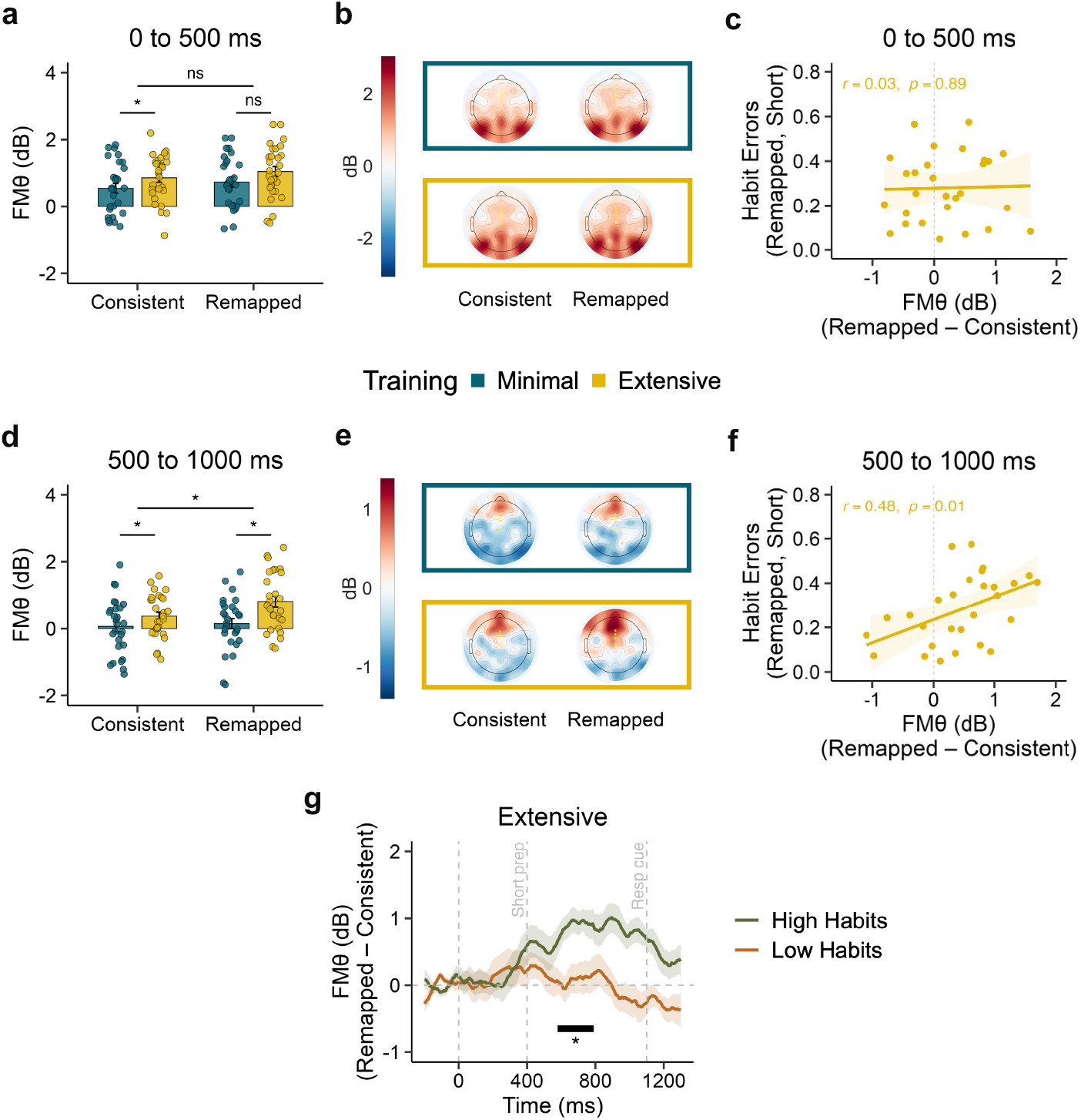
FMθ activity tracks reactive goal-directed control in high-habit individuals. **a**, FMθ power during the early window (0–500 ms) was higher on remapped than consistent trials (*F*(1,27) = 4.73, *p* = .039) and higher after extensive than minimal training (*F*(1,27) = 6.64, *p* = .016), with no interaction (*p* = .983). **b**, Early-window topographies show theta power by training condition and trial type. Yellow dots indicate midline electrode sites (FCz, Fz, Cz). **c**, FMθ contrast (remapped – consistent) in the early window was unrelated to habit errors after extensive training (*r* = .03, *p* = .890). d, FMθ power in the late window (500–1000 ms) showed a significant interaction (*F*(1,27) = 4.25, *p* = .049). **e**, Late-window topographies show elevated FMθ power on remapped trials after extensive training. **f**, In the late window, FMθ contrast after extensive training significantly predicted habit errors (*β* = 0.104, *p* = .010). **g**, After extensive training, participants with high habit errors (median split) showed significantly greater FMθ contrasts than those with low habit errors between 578 and 791 ms (cluster-based *p* = .033). Dots reflect individual participants. Bars and error bars show mean ± 1 SEM (a, d); shaded regions represent ± 1 SEM (g). \**p* < .05; \*\**p* < .01; ns = not significant.

We next tested whether FMθ power predicted individual differences in habit expression. If FMθ reflects the engagement of goal-directed control to counteract habitual response tendencies, then individuals with stronger habits should need to recruit more control to respond correctly. We regressed the proportion of habit errors on FMθ power from remapped trials after extensive training. Contrary to our prediction (H_5_), this association was not significant (*β* = 0.007, *SE* = 0.038, *t*(26) = 0.18, *p* = .861). One possibility is that baseline differences in theta power across individuals obscure a condition-specific effect. To account for this, we computed the FMθ contrast (remapped minus consistent), isolating the relative increase in control on trials where conflict between habitual and correct responses must be resolved. This was also not significant in the 0–500 ms window (*β* = 0.007, *SE* = 0.048, *t*(26) = 0.14, *p* = .890; Fig. 3c).

It is possible that goal-directed engagement unfolds later in the trial than we anticipated, particularly if control is recruited reactively following the initial detection of conflict, which we discovered occurred closer to 400 ms. To test this, we conducted an exploratory analysis on the later time window (500–1000 ms). Here, we found significant main effects of mapping (*F*(1,27) = 6.14, *p* = .020) and training (*F*(1,27) = 10.64, *p* = .003), with FMθ power higher on remapped than consistent trials and higher after extensive than minimal training. Notably, these factors also interacted (*F*(1,27) = 4.25, *p* = .049; Fig. 3d,e). Follow-up tests showed that the training effect was significant for both mappings but was more pronounced on remapped trials (*t*(27) = −3.37, *p* = .002) than consistent trials (*t*(27) = −2.14, *p* = .042). FMθ was significantly higher on remapped than consistent trials after extensive training (*t*(27) = −3.26, *p* = .003) but not after minimal training (*t*(27) = −0.60, *p* = .552). This pattern is consistent with greater reactive control engagement when extensively trained habitual responses must be overridden but should be interpreted with caution given the exploratory nature of the time window selection.

Given we found an interaction at this later time point, we explored whether individual differences in FMθ were related to habit expression in this time window also. Raw FMθ power on remapped trials after extensive training were not significantly associated with habit errors (*β* = 0.055, *SE* = 0.033, *t*(26) = 1.65, *p* = .111). However, the FMθ contrast revealed a significant association (*β* = 0.104, *SE* = 0.038, *t*(26) = 2.76, *p* = .010; Fig. 3f). Individuals with more habit errors showed larger FMθ in remapped versus consistent trials after extensive training. To characterize the precise timing of this relationship, we divided participants into high and low habit groups by median split on habit errors after extensive training and carried out a cluster-based permutation analysis on the FMθ contrast time course. This revealed that the high-habit group showed significantly greater FMθ contrasts than the low-habit group between 578 and 791 ms (cluster-based *p* = .033; Fig. 3g). Together with the LRP findings, this supports a dynamic model in which goal-directed control is flexibly engaged to counteract covert stimulus-driven activation when time allows. That said, it is important to acknowledge that we hypothesized these phenomena would occur earlier in the trial than they did.

### Extensive practice speeds up motor preparation but only LRP amplitude during training is associated with habit expression

We next examined how neural signatures of motor preparation and goal-directed engagement evolved across the course of extensive training. Analyses focused on correct trials from the training sessions on Day 1 and Day 14. Whereas the forced-response test captured competition between stimulus-driven and goal-directed processes, these analyses asked how training shapes the baseline dynamics of these systems when mappings remained stable and thus without trial-by-trial conflict.

Our preregistered analysis tested whether stimulus-locked LRP peak amplitude increased with extended training (H_6_), which would reflect a strengthening of stimulus-driven motor activation. Against our prediction, peak amplitude did not differ significantly between Day 1 and Day 14 (–3.58 ± 1.80 µV vs. –3.43 ± 1.64 µV; *t*_27_ = 0.52, *p* = 0.609; Fig. 4a), indicating that the overall strength of motor activation remained stable with training at the group level. However, although group-level amplitude did not change, participants who showed greater increases in peak LRP amplitude across training committed more habit errors, *r*(26) = 0.52, *p* = 0.004 (Fig. 4b), indicating that the degree to which training strengthened habitual motor activation predicted subsequent habit expression.

**Fig. 4:**
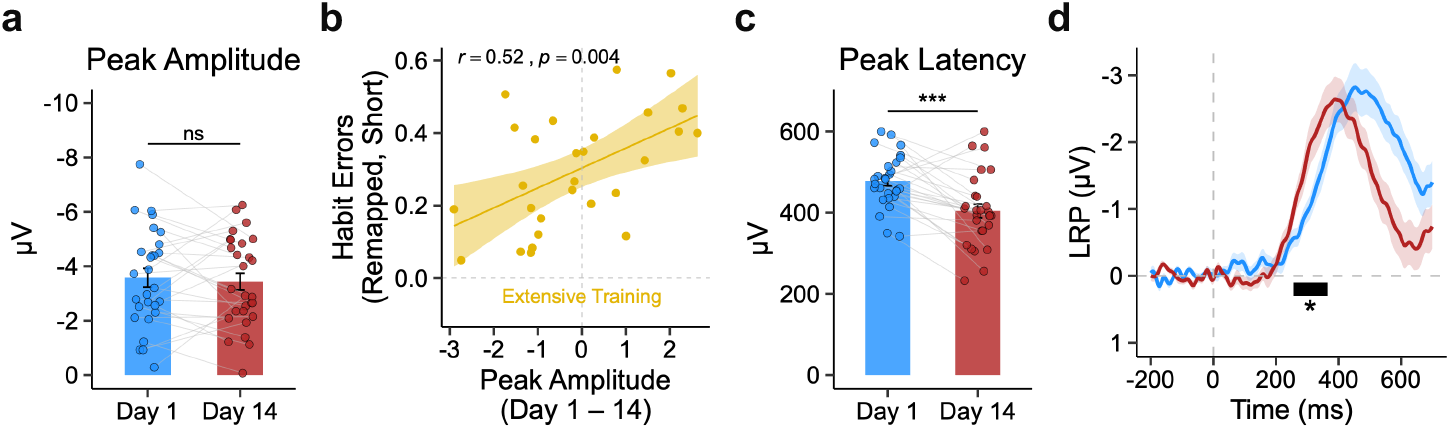
Extensive training decreases the latency of the LRP, but only its amplitude relates to individual differences in habit formation. **a**, Peak amplitude did not change with training (*t*_27_ = 0.52, *p* = 0.609). **b**, Increases in LRP amplitudes across training (i.e., more negative potentials, reflecting stronger motor activation) were associated with more habit errors for remapped, short trials after extensive training (*r* = 0.52, *p* = 0.004). **c**, Peak latency was significantly earlier on Day 14 compared to Day 1 (*t*_27_ = 4.47, *p* < 0.001), indicating faster motor preparation. **d**, Stimulus-locked LRP time series for Day 1 (blue) and Day 14 (red). A significant difference emerged between 256 to 365 ms (black bar; cluster-based *p* = 0.038).

Exploratory analyses revealed that training did affect the timing of motor preparation. Peak latencies of the LRP occurred earlier on Day 14 compared to Day 1 (404 ± 88 ms vs. 478 ± 64 ms; *t*_27_ = 4.47, *p* < 0.001; Fig. 4c), consistent with practice-related speeding of response preparation. A cluster-based permutation analysis on the stimulus-locked LRP (0 to 500 ms) confirmed that amplitudes for Day 1 and Day 14 significantly differed between 256 and 365 ms (cluster-based, *p* = 0.038; Fig. 4d), reflecting a faster signal build-up after extensive training. Reductions in LRP latency were, however, unrelated to habit expression, *r*(26) = -0.20, *p* = 0.303. Thus, while repetition promotes faster response preparation, the critical driver of individual differences in habitual behavior appears to be the strength, rather than the speed, of stimulus-driven response activation.

No differences were found for response-locked LRPs (peak amplitude: *t*_28_= –1.11, *p* = 0.275; peak latency: *t*_28_ = –0.48, *p* = 0.634), indicating that training influenced motor preparation but not execution.

We next examined stimulus-locked FMθ to test if goal-directed engagement reduces with extensive practice. Contrary to prediction (H_7_), FMθ did not differ between Day 1 and Day 14 (stimulus-locked: *t*_28_ = –0.89, *p* = 0.382; response-locked: *t*_28_ = 0.64, *p* = 0.525), indicating that goal-directed engagement remained stable over the course of training. The changes in FMθ from Day 1 to 14 were also not associated with habit errors (*β* = −0.019, *SE* = 0.033, *t*(27) = −0.57, *p* = .572).

Finally, we tested whether trait goal-directed capacity relates to individual differences in habit formation (H_8_). We reasoned that individuals with stronger goal-directed control might form stronger habits if attentional and control resources support more efficient encoding of S-R associations. To test this, we used model-based planning as a proxy for goal-directed capacity, measured via a separate two-step sequential decision-making task in which model-based planners prospectively evaluate action sequences to maximize reward^32^. Perhaps unsurprisingly, given the small *N*, individual differences in model-based planning were not associated with peak LRP amplitude on Day 14, (adjusting for baseline amplitude on Day 1) (*β* = –1.15, SE = 1.18, *p* = 0.34).

## Discussion

Our study provides a time-resolved account of how stimulus-driven and goal-directed systems compete during habit expression. We first replicated the behavioral finding that habits are expressed most strongly on short preparation trials, after an extended period of training. Then, we examined the LRP as a marker of motor preparation, finding that covert habit activation was detectable even on long response preparation trials (1100 ms) where participants successfully selected the correct action. Our preregistered analysis revealed a crossover pattern where extensive training enhanced motor preparation for the correct response on consistent trials while simultaneously increasing covert preparation of the inappropriate (now-incorrect) habitual response on remapped trials. Exploratory analyses localized this effect to a specific time window (379–479 ms), which was later than we had anticipated. Crucially, the strength of this covert activation was associated with habit expression on independent trials with short (400 ms) preparation time designed to reveal habit. These findings are consistent with a long-standing tenet of dual systems theories that habit strength is dissociable from habit expression under conditions where goal-directed control can be engaged.

Examining FMθ power as a marker of goal-directed control, our preregistered analysis in the 0–500 ms window revealed that FMθ was higher on remapped than consistent trials, likely reflecting greater recruitment of cognitive control to resolve conflict between habitual and correct responses. FMθ was also higher after extensive than minimal training, possibly reflecting increased control demands in anticipation of stronger habitual tendencies. However, the predicted interaction was not significant. Exploratory analyses in a later time window (500–1000 ms) did however reveal a significant interaction, with FMθ selectively elevated on remapped trials after extensive training. The later timing of this effect is consistent with the timing of habitual motor activation observed in the LRP (379–479 ms) and possibly points to goal-directed control being recruited reactively. Individuals who made more habit errors on short response preparation trials showed a greater selective increase in FMθ on remapped relative to consistent trials, suggesting that those with stronger habitual tendencies needed to recruit more cognitive control to respond correctly. These results are in line with formal models of arbitration under speed-accuracy trade-offs, which posit that fast but inflexible habitual policies are favored when response preparation time is scarce, whereas slower goal-directed selection dominates when deliberation is possible^33^. Though it should be noted these findings occurred later than we had predicted in our pre-registration.

This study sheds new light on not only habit expression, but also how extensive training causes habits to develop. Behaviorally, greater reaction time improvements across training were associated with stronger habit expression, suggesting that automatization through repeated practice drives habitual responding^34^. At a neural level, we observed faster LRP latencies after training, replicating prior work^35^, but this earlier onset was unrelated to individual differences in habit expression. This apparent dissociation could simply be because behavioral indicators are more reliable than neural ones. However, RTs reflect the outcome of numerous processes other than motor preparation including the speed of stimulus evaluation, response retrieval, and response execution, and it is thus unclear which component drove the association with habit expression. By examining the LRP, we could specifically isolate activation strength, indexed by peak amplitude, as the component related to individual differences in habit expression. Thus, while repetition promotes automatization, the critical driver of habitual behavior appears to be the strength of stimulus-driven response activation.

Most prior work has probed the mechanisms supporting habit expression by leveraging variation in goal-directed control. For example, showing that habit expression increases under stress and is linked to lower working memory capacity^36,37^ and reduced model-based planning^38–40^. This has meant that habit expression has been studied almost exclusively as failure of goal-directed control, with little understanding of the role of S–R associations^11,17^. This is because efforts to manipulate stimulus-response learning, for example through overtraining, have until recently been unsuccessful^17,18,41^. Research has shown that even 20 days of overtraining may not produce habits if individuals have sufficient time to suppress the stimulus-response system^19^. The time-dependent paradigm employed in the present study appears to surmount these challenges^19^. Here, we provide converging evidence consistent with dual systems accounts of habit expression. Our results suggest that habit expression may reflect a balance between habitual activation strength and the sufficiency of goal-directed control to override it. This could explain why some manipulations intended to impair goal-directed control, including stress, do not reliably lead to increased habit expression^42,43^. If the habitual activation itself is weak, impairing control may not be sufficient to produce habitual errors.

From a translational perspective, the careful separation of these neural systems may help identify distinct pathways conferring risk for repetitive behaviors across psychiatry. However, because achieving a balance between these systems is important for adaptive behavior, there are likely complex inter-dependencies and compensatory adaptations to consider. For example, recent evidence suggests that compulsive disorders may be associated with exaggerated S-R tendencies and impaired goal-directed control. Behavioral deficits in goal-directed control have been consistently linked to compulsivity^38,44^, in addition to reduced FMθ power during control-demanding tasks^45^. On the other hand, clinical studies of OCD have also revealed hyperactivation of motor preparation, with larger LRP amplitudes despite unchanged onset or latency^46,47^. More recently, there is behavioral evidence for accelerated habit formation under time pressure in compulsivity, using the same paradigm employed in the present study^28^. By capturing both covert habit activation and reactive control recruitment, our neural markers may provide a mechanistic bridge to understand how these systems function in compulsivity. More broadly, the ability to quantify habit strength and control engagement independently may provide a new avenue for developing markers to track vulnerability and evaluate treatment response in disorders of behavioral inflexibility.

Several limitations should be acknowledged. Although EEG affords excellent temporal resolution, its spatial precision is limited, and the LRP itself is a motor-preparation signal that we use to infer the strength of stimulus-driven activation. Combining EEG with Magnetoencephalography (MEG) or fMRI for source localization will be critical for linking these dynamics to corticostriatal circuits implicated in stimulus-driven and goal-directed control. Further, our task, while capturing core features of habit learning, was relatively simple and did not include an extrinsic reward structure. The relative stability of goal-directed engagement across training may therefore reflect the task’s low goal-directed demands, in contrast to real-world behaviors that unfold in complex environments involving conflict, context, and changing motivation. It is also important to acknowledge what the present data can and cannot establish. The preregistered LRP interaction was significant, providing confirmatory evidence for covert habitual activation. However, the preregistered brain-behavior analyses and the FMθ interaction were not significant in their preregistered forms – we had hypothesized timings of events consistently earlier than they occurred in the data. The significant brain-behavior effects emerged only in exploratory analyses using later time windows and in one case, a contrast measure. However, these findings were bolstered by convergent evidence across causal (i.e. training effects) and correlational analyses (relating EEG to habit expression) at same those time windows. Nonetheless, they warrant replication before strong conclusions can be drawn.

Questions have been raised about the construct validity of habits measured in laboratory tasks^21,48–50^. Researchers have struggled to find reliable correlations between different habit paradigms and as noted earlier, to consistently demonstrate habit expression as a function of training. Moreover, habit expression varies substantially across different tasks^21,29,50,51^. For instance, the contingency remapping used here produces more robust errors than classic outcome devaluation tests^52^. Additionally, some researchers have questioned whether habit errors reflect S-R associations at all, showing instead that they may also result from goal-directed processes operating on outdated information, such as the rapid retrieval of old action-outcome expectancies^30,31^. In a similar vein, others have suggested that habits result from strategic automatic action selection^53^. The current findings provide evidence for a neural marker of stimulus-driven response activation that is independent of habit expression and future work can expand this approach to assess the potential contribution of alternative mechanisms, such as automatically retrieved outcome information or strategic processes, in different task contexts.

In conclusion, we provide evidence that habitual motor responses are covertly activated even when overt behavior remains goal-directed, and that the strength of this covert activation is associated with individual differences in habit expression. Exploratory analyses further suggest that goal-directed control, indexed by FMθ, is selectively engaged to counteract habitual tendencies. By isolating EEG signatures of both stimulus-driven activation and goal-directed control, our findings offer evidence consistent with dual-systems models of habit-control arbitration. This approach bridges a long-standing gap between theories of habit formation and their behavioral manifestations, while offering a translational framework for probing maladaptive repetition and compulsivity across clinical conditions.

## Acknowledgements

CMG is supported by a European Research Council (ERC) starting grant (ERC-H2020-HABIT) and EKB is supported by funding from the Government of Ireland Postdoctoral Fellowship Programme (GOIPD/2023/1510).

## Contributions

**Conceptualization**: CMG; **Methodology**: CMG, SS; **Software**: SS; **Formal analysis**: EKB, SS, JPG, RGOC, CMG; **Investigation**: EKB, SS, AC, NSM, KRD, PR; **Resources**: CMG; **Data Curation**: EKB; **Writing – Original Draft**: EKB, CMG; **Writing – Review & Editing**: EKB, SS, JPG, AC, NSM, KRD, PR, RGOC, CMG; **Visualization**: EKB; **Supervision**: CMG, RGOC; **Project administration**: EKB, SS, CMG; **Funding acquisition**: CMG.

## Ethics declarations

This study has been granted ethical approval by the School of Psychology Ethics Committee, Trinity College Dublin (project code: SPREC012022-14).

## Competing interests

The authors declare no competing interests.

## Methods

### Participants and ethics

The preregistered sample size was 30 participants. Thirty-three individuals were recruited from the Trinity College Dublin community via the SONA participant pool, newsletters, online social media groups, and word of mouth. Inclusion criteria required participants to be aged between 18 and 30 years, right-handed, native English speakers (or equivalent fluency), and to have normal or corrected-to-normal color vision. Exclusion criteria included a personal or familial history of epilepsy, personal neurological illness, head injury, or unexplained loss of consciousness. All participants provided written informed consent and received €115 reimbursement distributed across the three in-person sessions. University students could alternatively choose to receive course credit. The study protocol was approved by the Trinity College Dublin Ethics Board (approval code: SPREC012022-14).

Thirty-three participants completed the experiment; three were excluded for producing fewer than 50% on-time responses in the forced-response test, resulting in a final behavioral sample of 30 participants (mean age = 21.3 years, SD = 3.1; 19 female, 9 male, 2 non-binary). For the forced-response test EEG analyses, valid trials were defined as those that were correct, on-time (within ±100 ms of the response cue), and free of EEG artefacts. For training EEG analyses, valid trials were defined as those that were correct, free of EEG artefacts, and with reaction times below 1000 ms. Trials in which participants did not commit a response were excluded from all analyses. Participants were then excluded from specific analyses if they did not meet minimum trial count thresholds after preprocessing. For the LRP analyses, participants were required to have at least 20 valid long-preparation trials per response hand in each mapping condition (i.e., consistent-left, consistent-right, remapped-left, remapped-right), as the LRP is computed as a contralateral–ipsilateral difference wave requiring stable averages per hand. Seven participants did not meet this criterion, yielding a final LRP sample of *n* = 23. For the FMθ analyses, participants were required to have at least 20 valid long-preparation trials in each mapping condition combined across response hands. Two participants did not meet this criterion, yielding a final FMθ sample of *n* = 28. For the training EEG analyses, one participant was excluded due to insufficient valid trials, and one further participant was excluded from stimulus-locked training peak analyses due to a positive peak amplitude (indicating no reliable LRP), yielding *n* = 29 for training analyses overall and *n* = 28 for stimulus-locked training LRP peak analyses.

### Habit Task

Participants completed a habit task^19^, designed to train four stimulus–response (S–R) associations under minimal and extensive training conditions in a within-subjects crossover design. The task was administered across three in-lab EEG sessions and twelve remote training sessions. The order of the training conditions was counterbalanced across participants (see Fig.1a for a schematic of the timeline procedure).

On each trial (see Fig. 1b), an abstract stimulus appeared above four on-screen boxes, each corresponding to one of four response keys (Q, W, O, P). Correct responses turned the corresponding box green; incorrect responses turned it red. Two non-overlapping sets of abstract stimuli were used for the two training conditions, counterbalanced across participants. S–R pairings were randomly assigned, and each mapping required responses using the left and right index and middle fingers.

In the minimal training condition, participants completed the task in a single in-lab session. They first learned an original mapping to criterion (i.e., five consecutive correct responses for each of the four stimuli; Fig. 1c). Immediately afterward, they completed a revised mapping to criterion, in which two of the four S–R associations were switched across hands (see Fig. 1d).

In the extensive training condition, participants underwent 14 days of training. On the first day (in-lab), they learned a different S–R mapping to criterion and completed three additional blocks of 100 trials each. Over the next twelve days, they completed seven 100-trial blocks each day remotely. On the final day (in-lab), they completed three more blocks, yielding a total of 9,000 training trials in the extensive condition. Following this training, participants performed a remapped version, in which two associations were again switched across hands.

During criterion training, there was no time limit for responses and participants learned the stimulus–response pairings through trial and error. During training blocks, participants responded at their own pace and were instructed to respond as quickly as possible.

After learning the revised mapping in both training conditions, participants completed a forced-response test. Habit expression was defined behaviorally as overt responses in line with the original, now incorrect, S–R mapping (see Fig.1d). Each trial began with a sequence of three tones (700 ms apart), and participants were instructed to synchronize their response with the third tone (Fig. 1e). The stimulus appeared either 400 ms (short preparation) or 1100 ms (long preparation) before the response cue (Fig. 1e). Correct and incorrect responses were followed by green and red feedback, respectively for 300 ms, and responses outside the permitted window (±100 ms) triggered a “too early!” or “too late!” message. The test included 10 blocks of 56 trials (560 total).

At the start of each in-lab session, participants completed practice blocks to familiarize themselves with the training and forced-response procedures. In the training practice block (40 trials), participants responded to highlighted fingers shown on a keyboard display. In the forced-response practice, they synchronized responses with the third tone. Practice continued until participants produced on-time responses (±100 ms) on at least 25 of the most recent 40 trials (minimum of 80 trials total).

### Additional measures

Participants completed the two-step reinforcement learning task at baseline^32^ to assess individual differences in goal-directed control. This task distinguishes model-based from model-free strategies by quantifying the influence of task structure on first-stage choices following rewards. Additionally, they filled out the Motor Tic, Obsessions and Compulsions, Vocal Tic Evaluation Survey (MOVES) and the Obsessive-Compulsive Inventory-Revised (OCI-R) at baseline, for exploratory correlations with EEG indices of stimulus-driven and goal-directed control and to obtain effect sizes for future sample size calculations.

### Data analysis

EEG preprocessing and analysis were conducted using custom MATLAB scripts based on EEGLAB^54^. Statistical analyses were performed in R (version 4.4.0). All hypothesis tests (H_1_-H_8_) were conducted as specified in our preregistration unless otherwise noted. Exploratory analyses are explicitly labelled as such throughout.

#### EEG Data Acquisition and Preprocessing

EEG data were recorded using a 128-channel BioSemi ActiveTwo system at a sampling rate of 1024 Hz and processed offline using EEGLAB^54^ in MATLAB (R2024a). Data were downsampled to 512 Hz and low pass filtered at 40 Hz. For both training and forced-response test data, preprocessing included linear detrending, segmentation of stimulus-locked epochs from –500 to 2000 ms relative to stimulus onset, and baseline correction to the 200 ms pre-stimulus window. Channels with extreme variance or high artefact counts were marked for interpolation, which was conducted separately for each block to account for transient signal degradation. On average, participants had *M* = 3.01 channels (*SD* = 4.54) interpolated per block during training and *M* = 1.29 channels (*SD* = 1.17) during test.

Epochs were excluded if (1) voltage on any scalp electrode exceeded ±100 µV, or (2) the bipolar VEOG signal exceeded ±200 µV (–200 ms to response). For forced-response test trials, epochs were additionally excluded if (3) the response was incorrect, or (4) the response was not on-time (outside ±100 ms of the response cue). For training trials, epochs were additionally excluded if (3) the response was incorrect, or (4) the reaction time exceeded 1000 ms. Trials in which participants did not commit a response were excluded from all analyses.

Participants in the LRP sample (*n* = 23) retained *M* = 170.7 (*SD* = 40.0) valid long trials per session out of 280 (*M* = 87.4 consistent, *M* = 83.3 remapped). Participants in the FMθ sample (*n* = 28) retained *M* = 159.4 (*SD* = 46.5) valid long trials per session out of 280 (*M*= 81.9 consistent, *M* = 77.6 remapped). During training, participants retained *M* = 257.50 trials (*SD* = 32.10) out of 300.

#### Lateralized Readiness Potential (LRP)

Stimulus-locked and response-locked lateralized readiness potentials (LRPs) were computed on a per-participant basis. For each trial, data from two motor-preparation electrodes (C3 and C4; BioSemi labels D19 and B22, channels 54 and 115) were extracted. During the forced-response test, stimulus-locked LRPs were calculated from epochs spanning –200 to 1100 ms relative to stimulus onset, baseline-corrected to the – 200 to 0 ms window. On each trial, a contralateral–ipsilateral difference wave was computed (C4 minus C3 for left-hand responses; C3 minus C4 for right-hand responses), yielding a time-resolved measure of motor preparation. These difference waves were averaged separately for left- and right-hand responses and then collapsed to a single waveform per participant. Further, covert habit activation was measured as the positive area under the LRP curve (AUC) from 150–300 ms, reflecting a Gratton dip. To capture motor activity aligned to action execution, response-locked LRPs were computed from a window extending 700 ms before to the response. Using the same contralateral– ipsilateral subtraction, we computed trial-level response-locked difference waves and averaged them for each response hand before collapsing across hands. The peak LRP for training analyses was determined by obtaining the maximum negative value for each participant in the stimulus-locked 200 to 600 ms time window.

#### Frontal Midline Theta (FMθ)

Frontal midline theta (FMθ) power was computed on a per-participant basis to index cognitive control and conflict monitoring. Data were extracted from FCz, Fz, and Cz electrodes (BioSemi labels A1, C21, C22, C23). Current source density (CSD) transformation was applied to the EEG data using the CSDToolbox^55^ with default parameters (*λ* = 1 × 10^−5^, *m* = 4), enhancing spatial resolution of the neural signals.

Stimulus-locked epochs (–200 to 1100 ms) were baseline-corrected to the –200 to 0 ms window. Data were band-pass filtered from 4 to 8 Hz and transformed using the Hilbert method to derive the analytic signal and compute instantaneous power. Power values were converted to decibels (dB) relative to the pre-stimulus baseline. FMθ power was averaged across the three electrodes to produce a single time course for each participant and condition. Mean power was computed for the preregistered 0–500 ms post-stimulus window. An exploratory analysis was also conducted on a later 500–1000 ms window, motivated by the possibility that goal-directed control is recruited reactively following initial conflict detection.

#### Brain–behavior analyses

To test whether EEG signatures predicted individual differences in habit expression (H_3_, H_5_), we regressed the proportion of habit errors (remapped, short preparation, extensive training) on the EEG measure of interest (LRP amplitude or FMθ power). The preregistration specified mixed-effects models for these analyses; however, because there was a single behavioral score per participant (proportion of habit errors after extensive training), these reduced to simple linear regressions. For LRP analyses (H_3_), the preregistered predictor was mean LRP amplitude from remapped trials after extensive training in the 150–800 ms window. An exploratory analysis also examined the 379–479 ms cluster window identified in the time-resolved analyses. For FMθ analyses (H_5_), the preregistered predictor was raw FMθ power on remapped trials after extensive training in the 0–500 ms window. Exploratory analyses additionally examined the FMθ contrast (remapped minus consistent) and the 500–1000 ms window.

#### Cluster-based permutation test

To assess time-resolved differences while controlling for multiple comparisons, we used cluster-based permutation tests using the *jlmerclusterperm* R package^56^. A significance threshold for clustering was set using the critical t-value from a two-tailed t-test (paired or unpaired depending on the comparison) at *α* = 0.05 with the corresponding degrees of freedom. Cluster-based permutation testing was performed with 5,000 permutations.

Clusters of contiguous time points exceeding the critical threshold were identified and p-values compared against *α* = 0.05. For 2 × 2 interaction analyses, difference scores were first computed to test for the predicted directional interaction effects. For between-group comparisons (median split analyses), unpaired t-values were used as the test statistic.

## Extended data

### Training performance

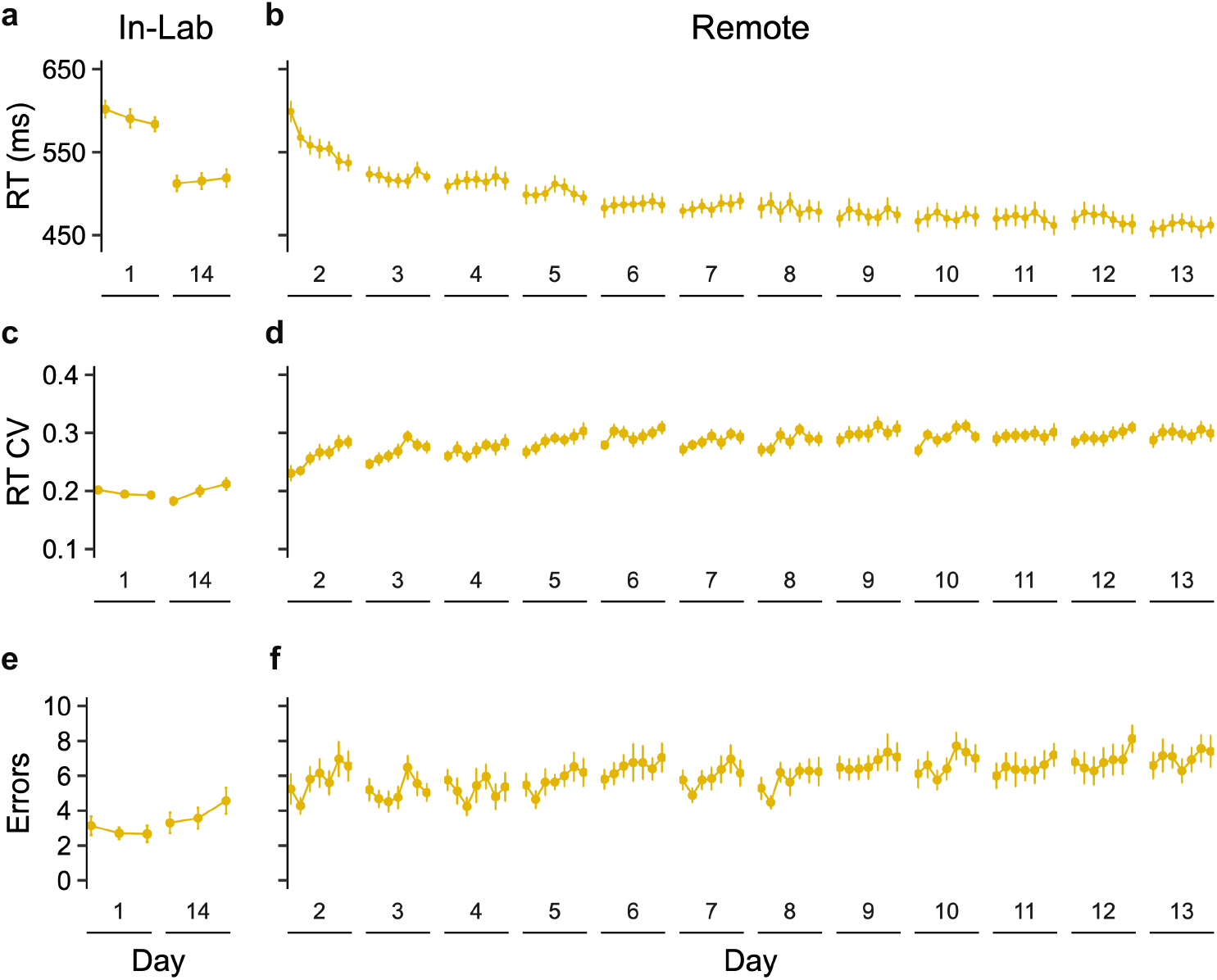
**a**, A significant interaction between Training Day and Block, *F*(66, 1584) = 3.10, *p* < .001. revealed significant within-day improvement on Day 2. **b**, For reaction time variability, there were significant main effects of Training Day, *F*(5.33, 128.00) = 7.02, *p* < .001, with lower variability on Day 2 compared to Day 14, and Block, *F*(3.53, 84.72) =12.45, *p* < .001, with lower variability in Block 1 compared to Block 7. **c**, Error rates were low and stable across blocks and days (all *F*s < 4.07). **d**, A repeated-measures ANOVA on RTs showed a significant interaction between Training Day and Block, *F*(66, 1584) = 3.10, *p* < .001. revealing significant within-day improvement on Day 2. **e**, For reaction time variability, there were significant main effects of Training Day, *F*(5.33, 128.00) = 7.02, *p* < .001, with lower variability on Day 2 compared to Day 13, and Block, *F*(3.53, 84.72) =12.45, *p* < .001, with lower variability in Block 1 compared to Block 7. **f**, For error rates, there were significant main effects of Training Day, *F*(5.52, 132.50) = 3.93, *p* = .002, and Block, *F* (3.89, 93.43) = 6.52, *p* < .001, showing higher error rates on Day 13 compared to Day 2, and in Block 7 than Block 1.

## Notes

https://osf.io/fm8q7/overview?view_only=5a02494967ad451eb6162dd566dcc8df

### Competing Interest Statement

The authors have declared no competing interest.

### Summary of Updates

Manuscript revised throughout in response to peer review.

